# Activity manipulation of an excitatory interneuron, during an embryonic critical period, alters network tuning of the *Drosophila* larval locomotor circuit

**DOI:** 10.1101/780221

**Authors:** Carlo N. G. Giachello, Yuen Ngan Fan, Matthias Landgraf, Richard A. Baines

## Abstract

As nervous systems develop, activity perturbations during critical periods can lead to permanently altered network function. However, how activity perturbation influences individual synapses, the network response and the underlying signalling mechanisms are not well understood. Here, we exploit a recently identified critical period in the development of the *Drosophila* larval locomotor circuit to show that activity perturbation differentially affects individual and identified synaptic pairings. Remarkably, we further show that activity-manipulation of a selective excitatory interneuron is sufficient to fully recapitulate the effects induced by network-wide activity disturbance; indicative that some neurons make a greater contribution to network tuning. We identify nitric oxide (NO)-signalling as a potential mediator of activity-dependent network tuning during the critical period. Significantly, the effect of NO-signalling to network tuning is dictated by the prior activity state of the network. Thus, this study provides mechanistic insight that is currently lacking into how activity during a critical period tunes a developing network.

## Introduction

Neural circuit development requires activity to refine connectivity, modify signalling properties and, ultimately, to ensure emergence of appropriate network output. As networks develop, sets of neurons become spontaneously active. Initial activity shows little coordination across the network. However, as circuits mature and, particularly, as sensory afferents become functional there is a shift towards increasingly coordinated patterned activity (1, 2). This transition is often marked by a critical period: a defined window of heightened plasticity during which activity exerts maximal influence to network properties (3). It is notable that errors made during a critical period can become ‘locked-in’ such that subsequently the network is unable to correct mistakes made during this period. A well described example is activity manipulation of visual inputs in developing mammals, which studies have shown have permanent and maladaptive effects to the formation of ocular dominance and can lead to clinical conditions such as amblyopia (4). While the existence of critical periods in developing circuits is well described and has revealed a number of important mechanisms, notably a role for a subset of GABAergic inhibitory neurons and the formation of perineuronal nets (3), a reliance on complex mammalian sensory circuits has precluded a cell-specific analysis of how activity, during a critical period, affects cell type-specific synaptic pairings and how such changes combine to impact network tuning.

We recently reported a critical period in the development of the *Drosophila* larval locomotor circuit, indicative that critical periods are a universal phenomenon. Manipulating activity late in embryogenesis (85-95% / 17 to 19h after egg laying) is sufficient to permanently alter the developmental trajectory of the locomotor network, leading to sub-optimal adjustment and instability. This manifests subsequently as a susceptibility to electroshock-induced seizures in larval stages, measured 5 days later at the end of larval life (5). Interestingly, the critical period coincides precisely with the emergence of patterned peristaltic contractions of body wall muscles in the developing embryo (6, 7). Appropriately patterned network activity is necessary for coordinated locomotor movements to emerge, indicative of this phase being essential for network tuning (8). Our previous work demonstrated that the overall level of network activity, during the critical period, is instructive for appropriate adjustment. For example, a gain-of function mutation in a voltage-gated Na^+^ channel (*para^bss^*) leads to increased activity levels and makes animals seizure prone. However, transient optogenetic reduction of activity during the critical period, in these mutants, is sufficient to allow appropriate tuning and entirely suppresses seizure susceptibility (5). At the single cell level, analysis of excitatory synaptic drive to motoneurons, both in *para^bss^* or control animals made seizure prone following activity manipulation during the critical period, reveals that excitatory currents have gained a significantly extended duration, suggesting a change to the excitation:inhibition balance (5).

The identification of a critical period in the *Drosophila* larval locomotor circuit presents a new opportunity to determine how synaptic transmission changes between specific, identified components of a network during normal tuning; and how these processes are impacted by transient activity manipulation during the critical period. One of the unique strengths of this locomotor network is that the upstream circuitry of segmentally re-iterated motoneuron-muscle pairs (~30 per half-segment (9)) has recently been characterised by tracing connectivity at the level of single synapses and functionally testing predicted connections (10–13). Thus, we are now able to identify and manipulate defined synaptic pairs and can ask, for the first time, how activity perturbation during a critical period alters signalling at identified synapses.

In this study, we focus on a synaptic pairing formed by an excitatory cholinergic premotor interneuron (termed ‘A27h’) and a postsynaptic motoneuron, ‘aCC’. The A27h interneuron is required for forward locomotion and likely contributes to intra-versus inter-segmental coordination of muscle contractions (12). In addition to innervating motoneurons (e.g. aCC) within the same segment, A27h also drives feed-forward inhibition via an inhibitory interneuron (GDL) in the next anterior segment, which in turn inhibits A27h in that same next anterior segment. This feed-forward inhibition has been proposed to facilitate the propagation of peristaltic-like contractions from posterior to anterior, which supports forward locomotion (12). We show that activity manipulation, during the critical period, affect synaptic pairings differently. For example, following transient network-wide activity manipulation during the critical period, overall cholinergic excitatory transmission onto the aCC motoneuron is increased, though the specific contribution by A27h is diminished. Remarkably, cell-specific optogenetic stimulation of the A27h interneuron during the critical period leads to a lasting increase in A27h→aCC synaptic transmission, suggestive of anti-homeostatic adjustments during this phase.

How networks achieve robust and reliable adjustment is not well understood, though hierarchical organisation or the presence of hub neurons have been suggested (14–16). Accordingly, we show that transient activity manipulation of the A27h interneuron is sufficient to disturb network development such that it fully recapitulates the effects of activity disturbance across the entire network. To understand the mechanistic basis for these effects, we implicate nitric oxide (NO)-signalling as mediating, at least in part, the effects of altered neuronal activity. Changing NO-signalling in the embryo mimics activity perturbations in that it is sufficient to make a network seizure prone. Moreover, the ability of increased activity during the critical period to alter synaptic strength and degrade network stability is blocked by NO inhibitors and potentiated by NO activators. We further show that, as with cell-selective activity changes, restricted manipulation of NO-signalling in A27h is sufficient to induce locomotor circuit instability, underlining NO-signalling as mediating effects of neuronal activity. Thus, we show that activity-manipulation during a critical period is sufficient to alter network tuning to produce a mature network that lacks robustness when challenged with a strong stimulus. Synapse-specific analysis shows that this is mediated by differential effects to identified synaptic pairings.

## Results

### Pan-neuronal activity perturbation during the critical period alters network excitation

We previously showed that transient perturbation of network activity during late embryonic development (17-19h after egg laying) leads to enduring network instability, measurable some five days later in third instar larvae (L3), as an increased seizure response following electroshock (Fig 1A). At the level of identified neurons, such transient activity manipulations during late embryogenesis, produces a characteristic broadening of excitatory cholinergic synaptic inputs (termed spontaneous rhythmic currents, SRCs) to identified motoneurons; in this instance the aCC motoneuron (Fig 1B) (5). SRCs are the product of the simultaneous activity of multiple premotor cholinergic interneurons (17). The same outcomes, to both increased seizure recovery time and SRC duration, are also caused by promoting hyperactivity in developing embryos by other means, e.g. exposure to the proconvulsant picrotoxin (PTX, a chloride channel blocker), or by introduction of the *para^bss^* mutation (a Nav hypermorph) (5). Equally, reducing synaptic excitation by inhibition during the critical period, e.g., by pan-neuronal activation of eNpHR, also leads to this effect. Therefore, changing activity levels in the developing network, to deviate from normal, irrespective of the nature or polarity of change, seems deterministic for, and detrimental to, post-embryonic network stability (5). Under conditions where the SRC inputs are broadened, we find the number of action potentials fired per SRC significantly increased (23.04 ± 3.32 *vs*. 12.08 ± 1.75 APs per bout, *para^bss^ vs*. wild-type, *p* = 0.004, unpaired *t*-test, Fig 1C-D), as measured by loose-patch extracellular recordings of endogenous spiking activity in aCC motoneurons, indicative of excessive excitation (see also (18)).

**Fig 1.**
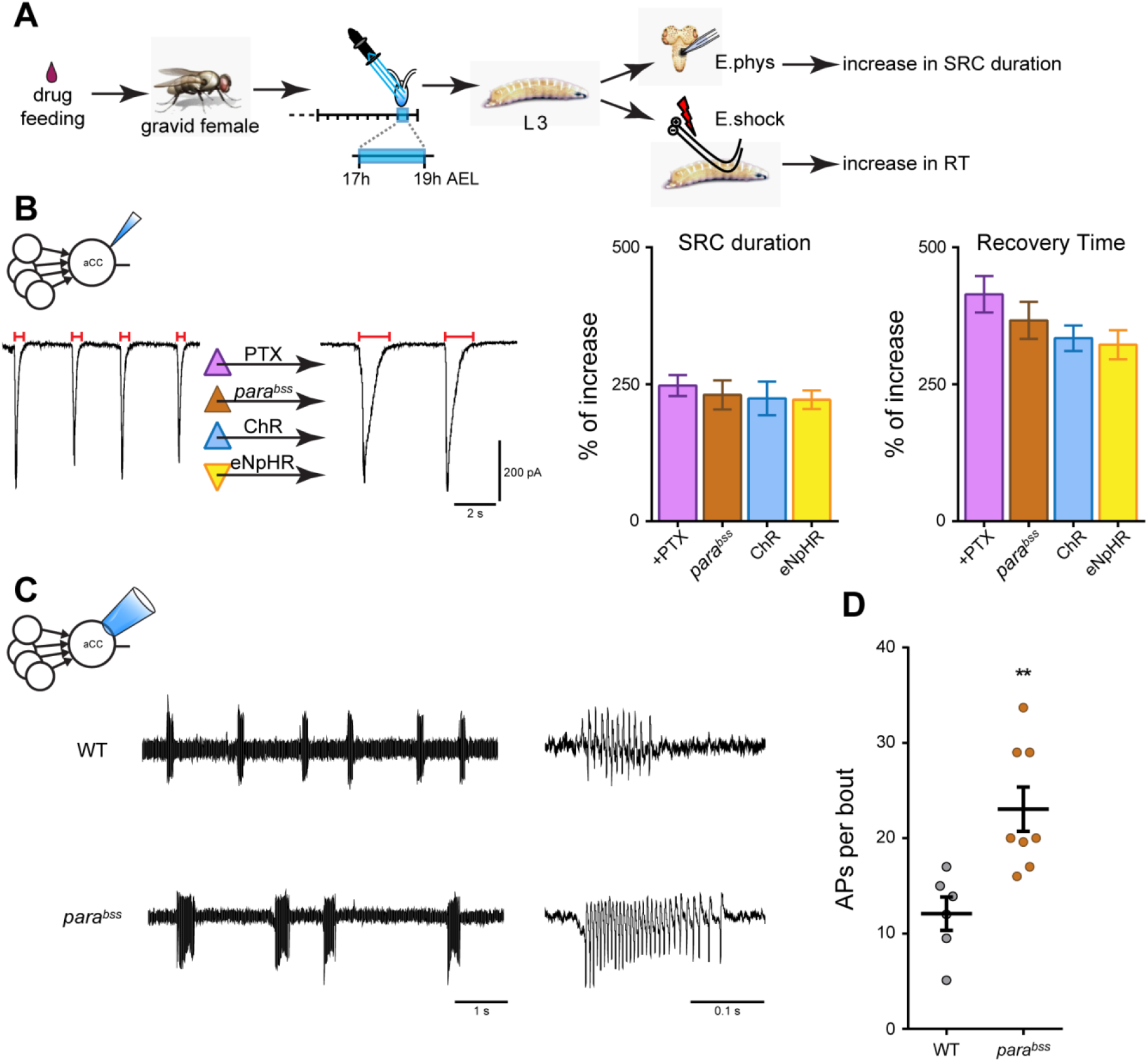
Activity manipulation during embryogenesis affects synaptic excitation of motoneurons and network stability. (A) Schematic representation of the experimental procedure. Early manipulation of neuronal activity was achieved by feeding drugs to gravid females (e.g. PTX). The drug is transferred to the embryo, but is no longer detectable at L3 (18). Alternatively, embryos expressing optogenetic constructs were exposed to light pulses (1 Hz) between 17-19h AEL (blue bar) which spans the identified critical period. 5 days later, L3 were tested by either electrophysiological recordings from aCC motoneurons or by electroshock, in order to measure change to synaptic drive or locomotor network stability, respectively. (B) Chemical (PTX), genetic (*para^bss^*), or optogenetic manipulation (*elav^C155^*>*ChR*, λ470 nm, 100 ms pulse duration/1 Hz; or, alternatively, *elav^C155^*>*eNpHR*, λ565 nm, 600 ms/1 Hz) during embryogenesis produces an increase in duration of SRCs recorded from L3 aCC (traces and left bar graph), which correlates with an increased recovery time to electroshock (right bar graph) (5). (C) Loose patch recordings from L3 aCC motoneurons showing increased endogenous spiking activity in conditions of excessive excitation of motoneurons (*para^bss^* mutation) compared to a wild-type strain (WT, Canton S). (D) Quantification of the number of action potentials per activity bout shown in panel C (*n* = 6 in WT, Canton S, and *n* = 8 in *para^bss^*, \*\**p* = 0.004). Data are represented as mean ± SEM.

### Neuron-specific activity perturbation during the critical period alters network excitation

The cellular mechanisms that ensure stable networks form and that might be affected by transient activity perturbation during a critical period, remain poorly understood. To study these, we took advantage of the fact that the larval *Drosophila* CNS is composed of identified neurons whose connectivity has been largely characterised (10–13). Specifically, we asked how activity manipulations of identified premotor interneurons, during the critical period, influence network function in later larvae. To test this idea, we focused on A27h: a cholinergic premotor interneuron that provides direct synaptic excitatory input to motoneurons, including the aCC motoneuron (Fig 2A) (12). We selectively manipulated activity of A27h, during the critical period, using *A27h-Gal4*>Chronos (100 ms/1Hz): a variant of ChR which exhibits a green-shifted excitation (λ565 nm, (19), see Methods for full rationale). Five days later, at the L3 stage, spontaneous SRC inputs to aCC (i.e. derived from multiple cholinergic pre-motor interneurons, not just A27h), showed an increased duration (0.98 ± 0.07 s *vs*. 0.46 ± 0.03 s, -LED *vs*. +LED, *p* < 0.0001, unpaired *t*-test, Fig 2B-C). In addition, electroshock of these L3 larvae showed a statistically significant increase in recovery time to 277 ± 16 s (+LED) from 83 ± 4 s (-LED, *p* < 0.0001, unpaired *t*-test, Fig 2D). Both outcomes are similar, if not identical, to changes caused following transient global network activity manipulations, see above (5).

**Fig 2.**
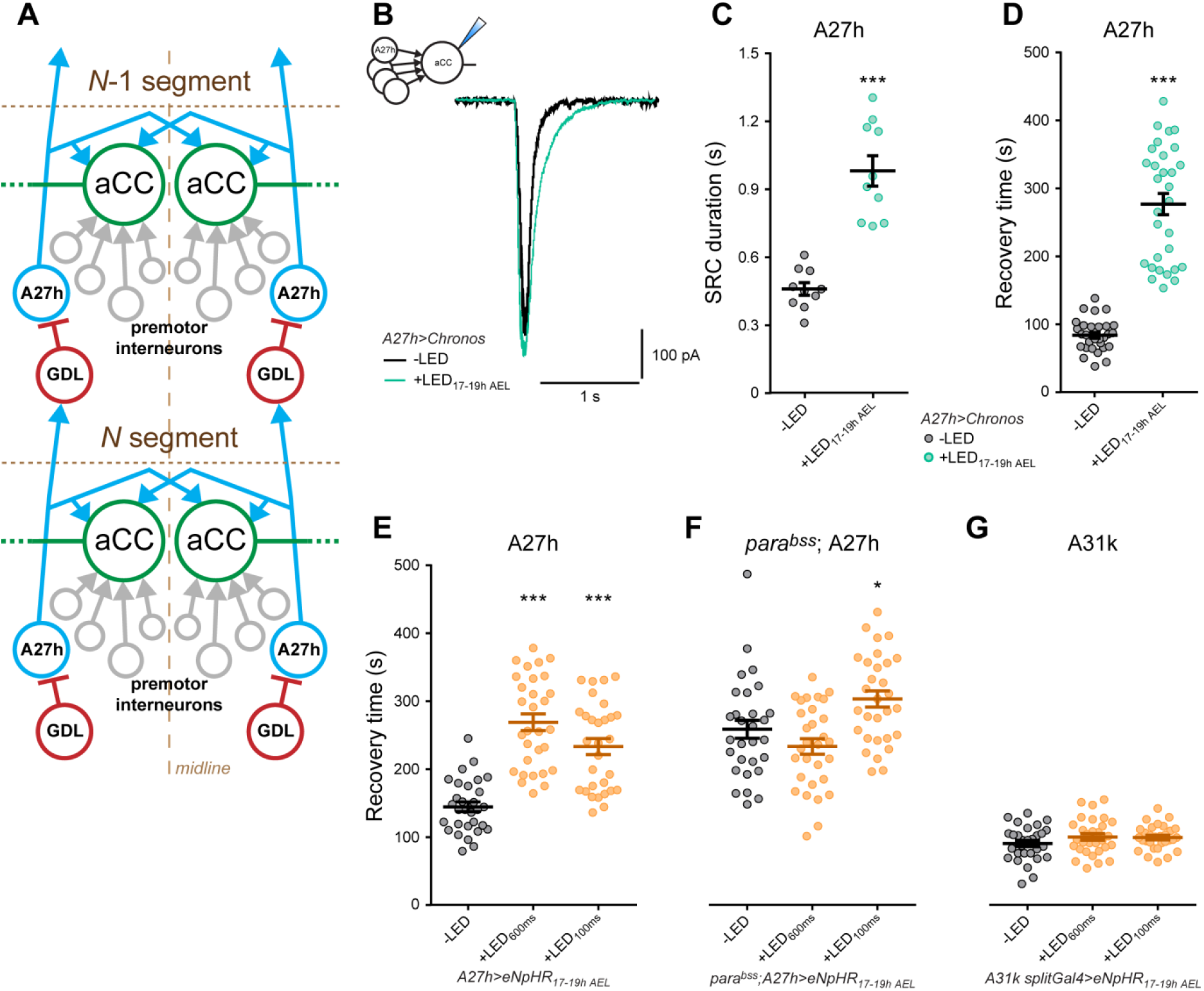
A27h manipulation produces network instability. (A) Schematic representation of the basic premotor circuit upstream of the aCC motoneuron. aCC receives inputs from several excitatory cholinergic interneurons, including A27h. A27h drives activity to aCCs located in the same segment (*N* segment), and also excites the GDL GABAergic interneuron, in the next anterior segment (*N*-1 segment). GDLs, in turn, inhibit the activity of A27h from the same segment, thus directing the propagation of the motor wave. Circuit adapted from (12). (B) Voltage-clamp recordings of endogenous (i.e. non-evoked) SRCs in L3 aCC motoneurons derived from embryos expressing Chronos in A27h and exposed to light (λ565 nm, 100 ms at 1Hz, 17-19h AEL). Transient activity manipulation of A27h, during the critical period, was sufficient to dramatically increase endogenous SRC duration. (C) Quantification of the SRC duration shown in panel B, *n* = 10 in each group, \*\*\**p* < 0.0001. (D) Analysis of recovery time from electroshocked L3 following identical embryonic activation of A27h, *n* = 30 in each group, \*\*\**p* < 0.0001. (E) Recovery time following electroshock of L3 derived from embryos expressing eNpHR in A27h and exposed to light (λ565 nm, 17-19h AEL). Longer light pulses (+LED_600 ms_) were used to suppress neuronal activity, while shorter pulses (+LED_100 ms_) produced hyperactivity due to rebound excitation (5). Both stimulations significantly increased recovery time compared to control (-LED). One-way ANOVA (F_(2, 87)_ = 36.25, *p* < 0.0001) followed by Bonferroni’s *post-hoc* test, *n* = 30 in each group, \*\*\**p* < 0.0001. (F) Inhibition of A27h (+LED_600 ms_, 17-19h AEL) was not sufficient to prevent network instability due to the presence of the *para^bss^* mutation. However, a significant additive effect of A27h excitation (+LED_100 ms_) and the *para^bss^* mutation was observed. One-way ANOVA (F_(2, 87)_ = 8.344, *p* = 0.0005) followed by Bonferroni’s *post-hoc* test, *n* = 30 in each group, \**p* = 0.035. (G) No detectable change in recovery time at L3 was observed following embryonic manipulation of A31k (λ565 nm, 100 and 600 ms, 17-19h AEL). One-way ANOVA (F_(2, 87)_ = 1.526, *p* = 0.223), *n* = 30 in each group. (C-G) Data are represented as mean ± SEM.

We obtained identical results by manipulating A27h with eNpHR. We previously demonstrated that eNpHR activation can generate two opposing effects dependent on light pulse duration: longer pulses (600 ms/1 Hz) produce sustained inhibition, while shorter pulses (100 ms/1 Hz) result in hyperactivity due to rebound firing, mimicking the effect of ChR or Chronos (5). Electroshock of L3 reared from manipulated embryos showed a statistically significant increase in recovery time following transient embryonic A27h inhibition (269 ± 12 *vs*. 144 ± 7 s, +LED_600ms_ *vs*. -LED, respectively, *p* < 0.0001, Fig 2E) or activation (233 ± 12 *vs*. 144 ± 7 s, +LED_100ms_ *vs*. -LED, respectively, *p* < 0.0001). We also observed that transient optogenetic activation of A27h (100 ms) potentiates the seizure severity caused by the *para^bss^* mutation (303 ± 12 *vs*. 258 ± 13 s, +LED_100ms_ *vs*. -LED, respectively, *p* = 0.035, Fig 2F). We previously showed that pan-neuronal inhibition during the critical period is sufficient to rescue the *para^bss^* seizure phenotype (5). However, NpHR-mediated inhibition of A27h (600 ms light pulses) was not sufficient to compensate for the *para*^bss^-induced network instability (233 ± 11 *vs*. 258 ± 13 s, +LED_600ms_ *vs*. -LED, respectively, *p* = 0.44). Thus, activity manipulation of A27h interneurons is sufficient to destabilise the locomotor network, similar in phenotype to global network activity manipulations; though in the context of an over-excitable network (e.g. in *para^bss^* mutants) selective inhibition of A27h activity is insufficient to prevent the network from becoming unstable.

Next, we asked whether manipulating the activity of any neuron during the critical period would be sufficient to cause a subsequent seizure recovery phenotype. We tested this idea by identical activity manipulations of the GABAergic interneuron A31k (*20A03-AD;87H09-DBD* split Gal4 line), which delivers proprioceptive feedback to motoneurons (11). We found that these did not produce any change in recovery time to electroshock treatment at L3 (Fig 2G). We confirmed that all lines express Gal4 during the embryonic period of optogenetic manipulation, evidenced by expression of fluorescent UAS-reporters (see Methods for details).

Combined, our results suggest that different interneurons may have varying propensities for driving change in network adjustment during the critical period, possibly with activity manipulations of excitatory, cholinergic neurons, being more contributory than manipulations of inhibitory GABAergic cells.

### Activity-manipulation differentially influences A27h synaptic drive

A key question is how critical period activity manipulations impact on synaptic transmission between partner neurons. We took advantage of having access to known partner neurons in this network and measured the output of A27h→aCC pairing, using optogenetics to selectively activate the A27h interneuron when recording its synaptic drive to the aCC motoneuron.

In controls, not exposed to embryonic manipulation, optogenetic stimulation of A27h (Chronos, λ565 nm, 1 s) produced an inward current in aCC (Fig 3A). Current density reached a maximum of 5.07 ± 0.66 pA/pF (*n* = 10, Fig 3B). Selective activity perturbation of A27h (A27h>Chronos, 100 ms/1Hz), during the critical period, resulted in a markedly increased A27h→aCC synaptic current amplitude recorded at L3 (11.27 ± 0.92 pA/pF, *n* = 10, *p* < 0.0001, Fig 3A-B). We then compared this with transient pan-neuronal activity manipulations during the critical period. For this we exposed embryonic networks to the proconvulsant picrotoxin (PTX) by feeding it to gravid females (using the GAL4 expression system to selectively stimulate A27h for recording at L3). We previously demonstrated that feeding drugs to gravid females leads to significant transfer of drug to the embryo. This is transient because by late larval stages, when our measurements are made, radio-tracing showed the drug is eliminated (18). Surprisingly, although embryonic exposure to PTX leads to an increase in overall synaptic drive to the aCC motoneuron (cf. Fig 1B), the specific A27h→aCC pairing synaptic transmission is significantly reduced (2.05 ± 0.22 pA/pF, *n* = 10, *p* = 0.0048, Fig 3A-B). We find the same reduction in A27h→aCC synaptic drive following a different global activity manipulation that similarly causes network instability, namely the *para^bss^* mutation (1.61 ± 0.19 pA/pF, *n* = 10, *p* = 0.001, Fig 3A-B).

**Fig 3.**
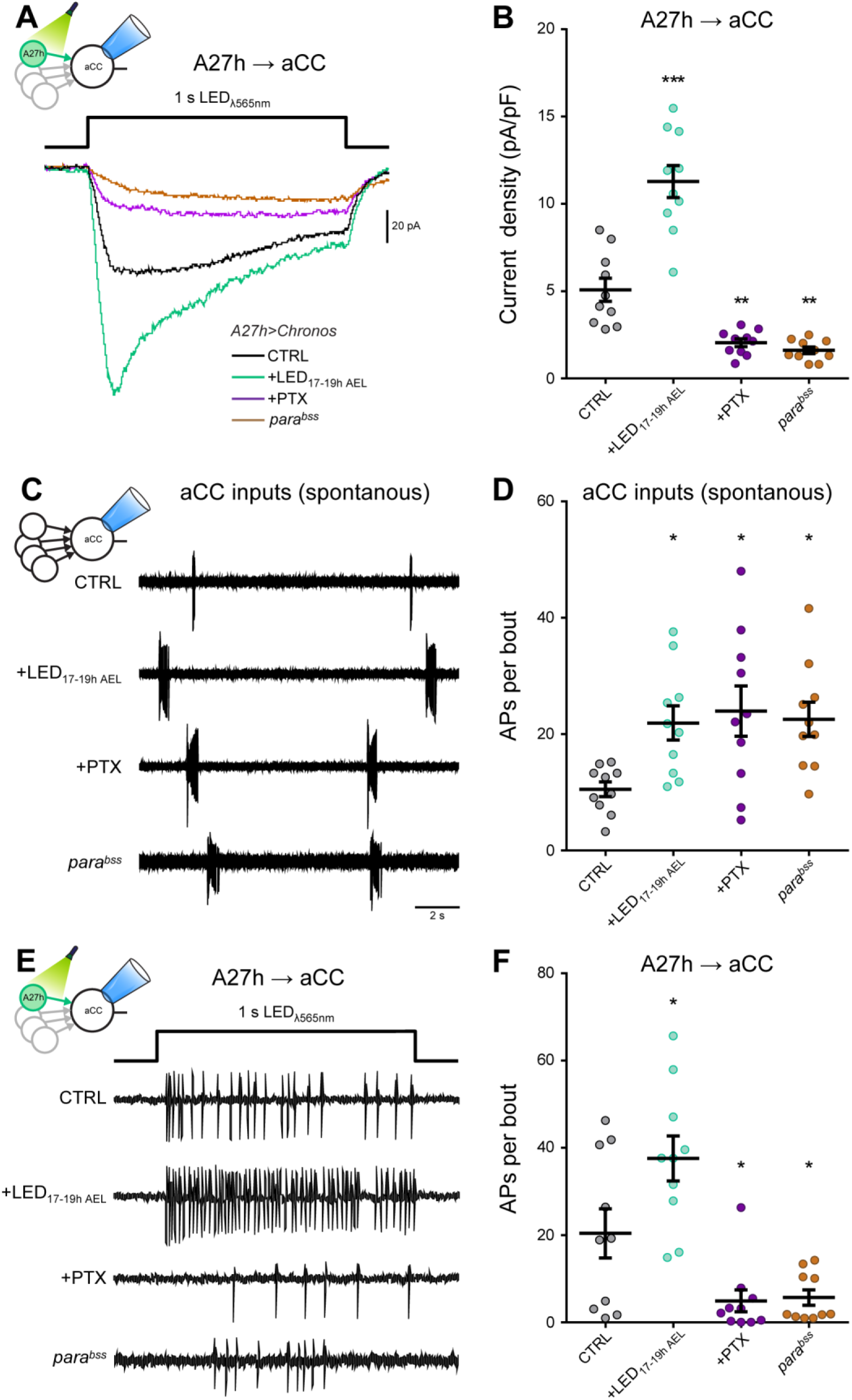
Pan-neuronal and A27h-specific activity perturbation differentially affects A27h drive to aCC. (A) Voltage-clamp recordings from L3 aCC showing the postsynaptic response to Chronos-induced stimulation of A27h (*A27h-Gal4*>*Chronos*, λ565 nm, 1s, black trace). Optogenetic-induced hyperactivity of A27h, during the critical period, is sufficient to potentiate A27h→aCC connectivity (+LED_17-19h AEL_, green trace). Conversely, global CNS hyperactivity, achieved by embryonic exposure to PTX or the *para^bss^* mutation, is sufficient to reduce A27h→aCC synaptic current amplitude (purple/brown traces). (B) Quantitative analysis of the synaptic drive from A27h→aCC. One-way ANOVA (F_(3, 36)_ = 58.16, *p* < 0.0001) followed by Bonferroni’s *post-hoc* test (*n* = 10 in each group, \*\*\**p* < 0.001, \*\**p* = 0.0048 and 0.001, respectively). (C) Loose patch recordings from L3 aCC motoneurons showing enhanced endogenous (spontaneous) spiking activity in the three experimental conditions compared to CTRL (*A27h-Gal4*>*Chronos*, unstimulated). (D) Quantification of the number of action potentials per bout shown in panel C. One-way ANOVA (F_(3, 36)_ = 4.099, *p* = 0.0133) followed by Bonferroni’s *post-hoc* test (*n* = 10 in each group, \**p* = 0.0375, 0.0112 and 0.0261, respectively). (E) Loose patch recordings from L3 aCC motoneurons showing the contribution of A27h (*A27h-Gal4*>*Chronos*, λ565 nm, 1s) to aCC spiking activity. Enhanced firing was observed following selective embryonic hyperactivity of A27h (+LED_17-19h AEL_). Conversely, fewer action potentials were measured after global CNS hyperactivity (+PTX and *para^bss^*). (F) Quantification of the number of evoked action potentials per bout shown in panel C. One-way ANOVA (F_(3, 36)_ = 13.94, *p* < 0.0001) followed by Bonferroni’s *post-hoc* test (*n* = 10 in each group, \**p* = 0.0166, 0.0349 and 0.0481, respectively). (B, D, F) Data are represented as mean ± SEM.

Under all three experimental conditions, loose patch extracellular recordings from L3 aCC motoneurons showed enhanced endogenous spiking activity (i.e. spontaneous activation of several premotor interneurons) (+LED: 21.92 ± 2.95, *p* = 0.0375; +PTX: 23.97 ± 4.31, *p* = 0.0112; *para^bss^*: 22.55 ± 2.95, *p* = 0.0261; *vs*. CTRL: 10.53 ± 1.26 APs per bout, Fig 3C-D). This suggests that both pan-neuronal and A27h-specific activity perturbations lead to excessive excitation of motoneurons, by increasing spontaneous SRC duration (see above) which, in turn, compromises network stability. In order to evaluate the specific contribution of A27h activation to aCC firing, we recorded aCC activity in loose patch mode while simultaneously optogenetically stimulating A27h (A27h>Chronos, λ565 nm 1 s pulses, Fig 3E). We observed that the increased synaptic drive resulting from the cell-selective embryonic manipulation of A27h (cf. Fig3A) is sufficient to make aCC fire more action potentials (from 20.40 ± 5.62 to 37.58 ± 5.17 APs per bout, *p* = 0.0166, CTRL *vs*. +LED, respectively, Fig 3E-F). In contrast, following transient global CNS perturbation during the critical period, the number of action potentials evoked by A27h activation is significantly lower (+PTX: 4.93 ± 2.51, *p* = 0.0349; *para^bss^*: 5.69 ± 1.77 APs per bout, *p* = 0.0481, Fig 3E-F).

Thus, we show that both global and cell-specific activity manipulations have comparable effects on network stability, evidenced by increased recovery times following electroshock-induced seizures, as well as endogenous aCC spiking activity. Using a cell-selective A27h manipulation, we show that at least for this excitatory interneuron, increased activation during the critical period leads to strengthened synaptic transmission for both this and other premotor cholinergic interneurons. In contrast, when network activity is globally manipulated during the critical period, transmission at the A27h→aCC pairing is reduced, but the excitatory drive to aCC from other premotor interneurons is increased which results is a similar outcome: network instability to electroshock. Thus, any deviation away from the normal strength of the A27h→aCC coupling, during the critical period, is sufficient to alter network tuning.

### Nitric oxide mediates activity perturbation during the critical period

The mechanisms underlying network tuning during a critical period are not well understood. Taking a best-candidate approach, we focussed on nitric oxide (NO)-signalling, since this messenger has been shown to be regulated by activity and able to alter synaptic drive (20). In a first set of experiments, we pharmacologically manipulated NO synthase (NOS) activity in embryos by feeding gravid females either the NOS inhibitor, N(G)-nitro-L-arginine methyl ester (L-NAME, 0.1 M), or the NO donor, sodium nitroprusside (SNP, 1.5 mM). On their own, these manipulations did not lead to changes in SRCs recorded from L3 aCC motoneurons or recovery times following electroshock-induced seizures (Fig. 4). We then asked whether SRC and seizure recovery phenotypes caused by activity manipulations during the critical period were mediated by NO-signalling. To test this hypothesis, we conducted the same pharmacological NO synthase manipulations in embryos, but this time additionally carried out optogenetic pan-neuronal activity manipulation (*elav^C155^*>*ChR*) during the critical period. Recordings from aCC in control L3 (absence of NO manipulation) showed an expected increase in SRC duration following optogenetic manipulation (0.52 ± 0.04 *vs*. 1.25 ± 0.13 s, -LED *vs*. +LED, respectively, *p* = 0.003, Fig 4A-B). The presence of the NOS inhibitor L-NAME (0.1 M) abolished SRC broadening (0.57 ± 0.03 *vs*. 0.61 ± 0.05 s, -LED *vs*. +LED, respectively, *p* > 0.9). Conversely, embryos from females fed the NO donor, SNP (1.5 mM), exhibited a significantly larger increase in SRC duration (0.49 ± 0.05 to 2.01 ± 0.27 s, -LED *vs*. +LED, respectively, *p* < 0.001). A two-way ANOVA showed both drug treatments produced significant changes in the optogenetically manipulated groups (*p* = 0.0148 and *p* = 0.0016, L-NAME and SNP *vs*. CTRL, respectively), whilst drug treatment per se had no impact on SRCs in non-stimulated (-LED) animals. Throughout, SRC frequency and amplitude were unaffected.

**Fig 4.**
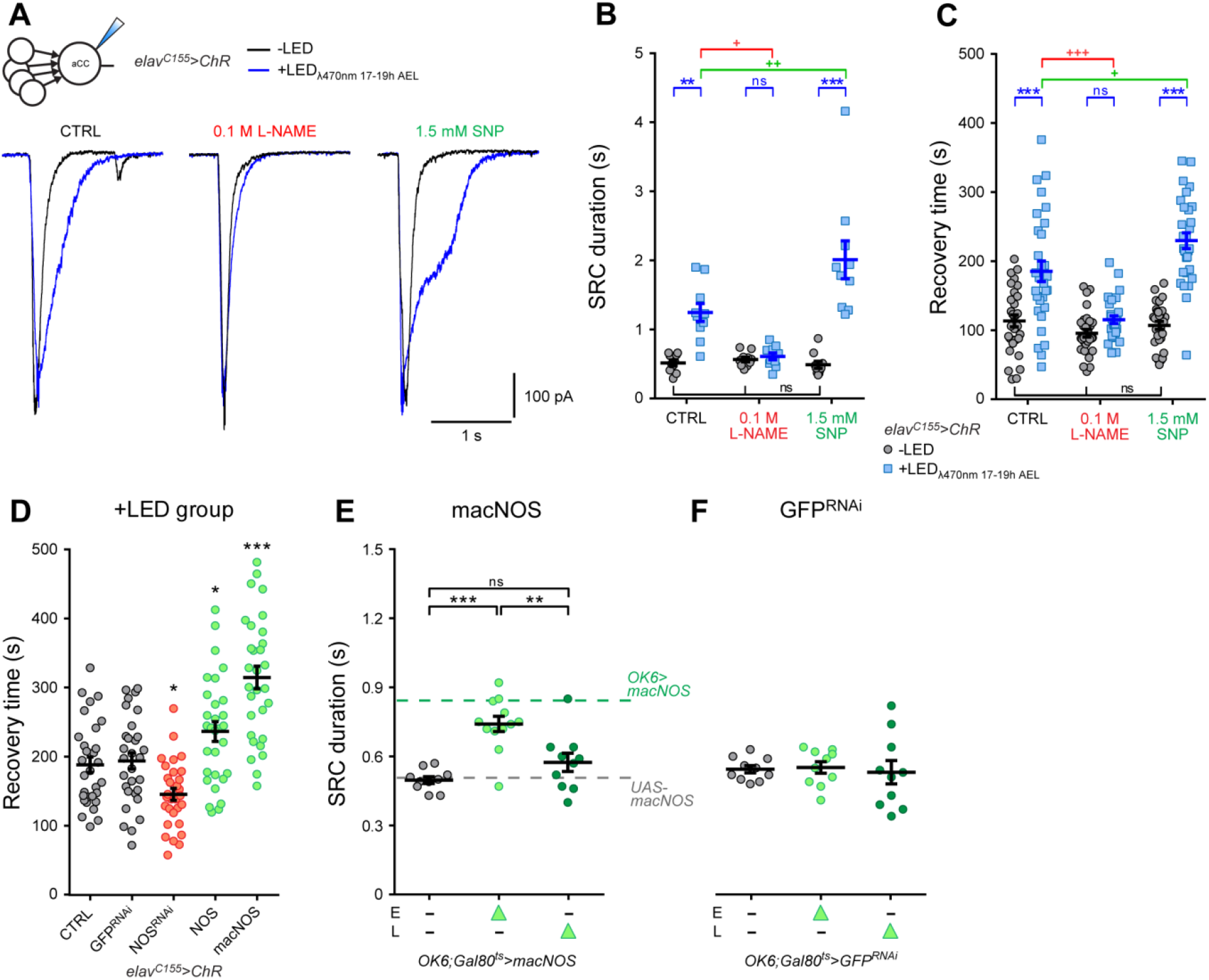
Nitric oxide mediates activity perturbation during the critical period. (A) Voltage-clamp recordings from L3 aCC show an increase in SRC duration following pan-neuronal activation of ChR during the critical period (*elav^C115^*>*ChR*: +LED 17-19h AEL, λ470 nm, blue trace, *vs*. −LED, black trace). Inhibiting NOS (0.1 M L-NAME), prior to optogenetic manipulation, blocks this effect, while exposure to the NO donor (1.5 mM SNP) potentiates SRC broadening. (B) Quantitative analysis of SRC duration, *n* = 10 in each group. A two-way ANOVA analysis revealed a significant effect of the +LED treatment (F_(1, 54)_ = 52.63, *p* < 0.001), NOS manipulation (F_(2, 54)_ = 13.16, *p* < 0.001), and interaction (F_(2, 54)_ = 16.4, *p* < 0.001). (C) Recovery time to electroshock measured at L3 following the same stimulation protocol. Exposure to the NOS-inhibitor L-NAME blocked the ChR-induced increase in recovery time, while feeding the NO donor SNP potentiated this effect even further, *n* = 30 in each group. A two-way ANOVA analysis revealed a significant effect of the LED treatment (F_(1,174)_ = 86.43, *p* < 0.001), NOS manipulation (F_(2, 174)_ = 23.54, *p* < 0.001), and interaction (F_(2, 174)_ = 15.1, *p* < 0.001). (B, C) \*\**p* < 0.01 and \*\*\**p* < 0.001 are significant to +LED *vs*. -LED within each group; ^+^*p* < 0.05, ^++^*p* < 0.01 and ^+++^*p* < 0.001 show significance to NOS drugs (+LED groups *vs*. CTRL), Bonferroni’s *post-hoc* test. (D) Recovery time from electroshock of L3 pan-neuronally co-expressing various transgenes together with ChR. Embryos were exposed to light (+LED group: λ470 nm, 100 ms, 17-19h AEL). NOS inhibition (NOS^RNAi^) reduced the expected ChR-induced increase in recovery time. By contrast, up-regulation of NOS-signalling (NOS and macNOS) potentiated the effect of ChR activation. Co-expression of GFP^RNAi^, as a control showed no change compared to CTRL. One-way ANOVA (F_(4, 145)_ = 25.38, *p* < 0.001) followed by Bonferroni’s *post-hoc* test, *n* = 30 in each group, \**p* = 0.0312 and 0.0324, respectively, \*\*\**p* < 0.001. (E) Temporal regulation of macNOS expression in motoneurons was achieved through GAL80^ts^. GAL4-mediated expression of macNOS during embryogenesis (n = 12) but not during larval stages (n = 10) led to an increase in SRC duration at L3. One-way ANOVA (F_(3, 29)_ = 16.36, *p* < 0.001) followed by Bonferroni’s *post-hoc* test, \*\*\**p* < 0.001. Dotted lines represent reference values obtained from *OK6*>*macNOS* and, as a further control, the *UAS-macNOS* parental line. (F) In identical temperature-controlled experiments, the expression of GFP^RNAi^, used as an additional control, did not show detectable change in SRC kinetics (*n* = 10 in each group). (B-F) Data are represented as mean ± SEM.

Embryonic exposure to the NOS-inhibitor, L-NAME (0.1 M), also counteracted electroshock-induced network instability by preventing the increased recovery time normally associated with pan-neuronal optogenetic manipulation (96 ± 6 *vs*. 115 ± 6 s, -LED *vs*. +LED, respectively, *p* > 0.9, Fig 4C). Conversely, embryonic exposure to the NO donor, SNP (1.5 mM), significantly potentiated the effects of optogenetic activity-perturbation, leading to a further increase in recovery time (from 107 ± 6 s to 230 ± 11 s, *p* < 0.001). A two-way ANOVA showed embryonic manipulation of NO levels significantly affected only the activity manipulated (+LED) groups (*p* < 0.0001 and *p* = 0.0156, L-NAME and SNP *vs*. CTRL, respectively), indicating that the concentrations of NOS modifiers used did not by themselves alter motor circuitry development. To further confirm the role of NO-signalling downstream of neuronal activity, we also tested the ability of additional drugs, known to target NOS and other members of the canonical NO-signalling pathway, including the soluble guanylyl cyclase receptor (sGC) and PKG kinase. Consistently, inhibitors of NO-signalling prevented increases in recovery time, while activators caused further potentiation (Fig S1).

We also targeted genetic manipulations of NO-signalling to developing neurons to validate pharmacology. Mirroring the effect of the NOS-inhibitor, L-NAME, pan-neuronal expression of NOS^RNAi^ was sufficient to block an increase of recovery time normally caused by optogenetic network activation during the critical period (*elav^C155^*>*ChR;NOS^RNAi^*: 145 ± 9 s, *p* = 0.0312, Fig 4D). Conversely, expression of *Drosophila* NOS (mimicking the activity of SNP) resulted in a potentiation (*elav^C155^*>*ChR;NOS*: 237 ± 14 s, *p* = 0.0324). For both manipulations, significant changes in *dNOS* mRNA, albeit in opposite directions, was confirmed by qRT-PCR (see Methods). Moreover, expression of a constitutively active transgene (*UAS-macNOS*), resulted in an even stronger potentiation of recovery time (*elav^C155^*>*ChR;macNOS*: 315 ± 16 s, *p* < 0.001), indicative of a dose-response. Co-expression of GFP^RNAi^ served as an additional control (*elav^C155^*>*ChR;GFP^RNAi^*) and showed no change compared to CTRL (194 ± 12 *vs*. 189 ± 11 s, GFP^RNAi^ *vs*. CTRL, respectively, *p* > 0.9).

Whilst our previous work has shown that drugs fed to gravid females enter the embryo, perdurance through to final third larval instar does not occur (Marley and Baines, 2011). However, a caveat to results obtained using a Gal4 driver is its expression throughout both embryonic and larval periods. To show that embryonic, but not larval, manipulation of NO-signalling is sufficient to alter synaptic drive we manipulated Gal4 activity with a temperature sensitive Gal4 inhibitor, Gal80^ts^ (21). Restricting expression of macNOS to motoneurons only during embryogenesis (*OK6;Gal80^ts^*>*macNOS*) resulted in a significant increase in SRC duration in aCC recorded later in L3 (from 0.50 ± 0.01 s to 0.74 ± 0.03 s, *n* = 12, *p* < 0.001, Fig 4E). By contrast, no change in SRC duration was observed when expression of macNOS was restricted to postembryonic larval stages (0.57 ± 0.04 s, *n* = 10, *p* = 0.31). To verify that the temperature changes required for GAL80^ts^ did not affect SRC kinetics, the same temperature shifts were repeated using a control genotype (*OK6;Gal80^ts^*>*GFP^RNAi^*), which showed no significant effects to SRC kinetics (Fig 4F).

### Altered NO-signalling in A27h recapitulates changes caused by activity manipulations

Our results support the hypothesis that activity regulated changes, during network tuning, are mediated, at least in part, through NO-signalling. If this is indeed the case, and if NO-signalling plays a major role in network adjustment during the critical period, then one might expect manipulations of NO-signalling restricted to neurons such as A27h to mimic effects of cell-specific activity manipulations. To test this, we knocked down NOS transcript (*A27h*>*NOS^RNAi^*) or increased NO-signalling (*A27h*> *UAS-macNOS*) in A27h. As predicted, this led to a significantly increased recovery time to electroshock in both conditions (*A27h*>*NOS^RNAi^*: 172 ± 10 s, *p* < 0.0001; *A27h*>*macNOS*: 188 ± 10 s, *p* < 0.0001, *vs*. *A27h*>*GFP^RNAi^* control: 103 ± 6 s, One-way ANOVA, Fig 5A). As a further control, we expressed the same constructs in the GABAergic interneuron A31k, where activity manipulation was ineffective (cf. Fig 2G). No significant change was detectable following NOS manipulation (*A31k*>*NOS^RNAi^*: 122 ± 5 s, *p* = 0.1744; *A31k*>*macNOS*: 94 ± 5 s,*p* = 0.1007, *vs*. control: 109 ± 5 s, One-way ANOVA, *n* = 30 in each group), suggesting a crucial role for A27h in network stability.

**Fig 5.**
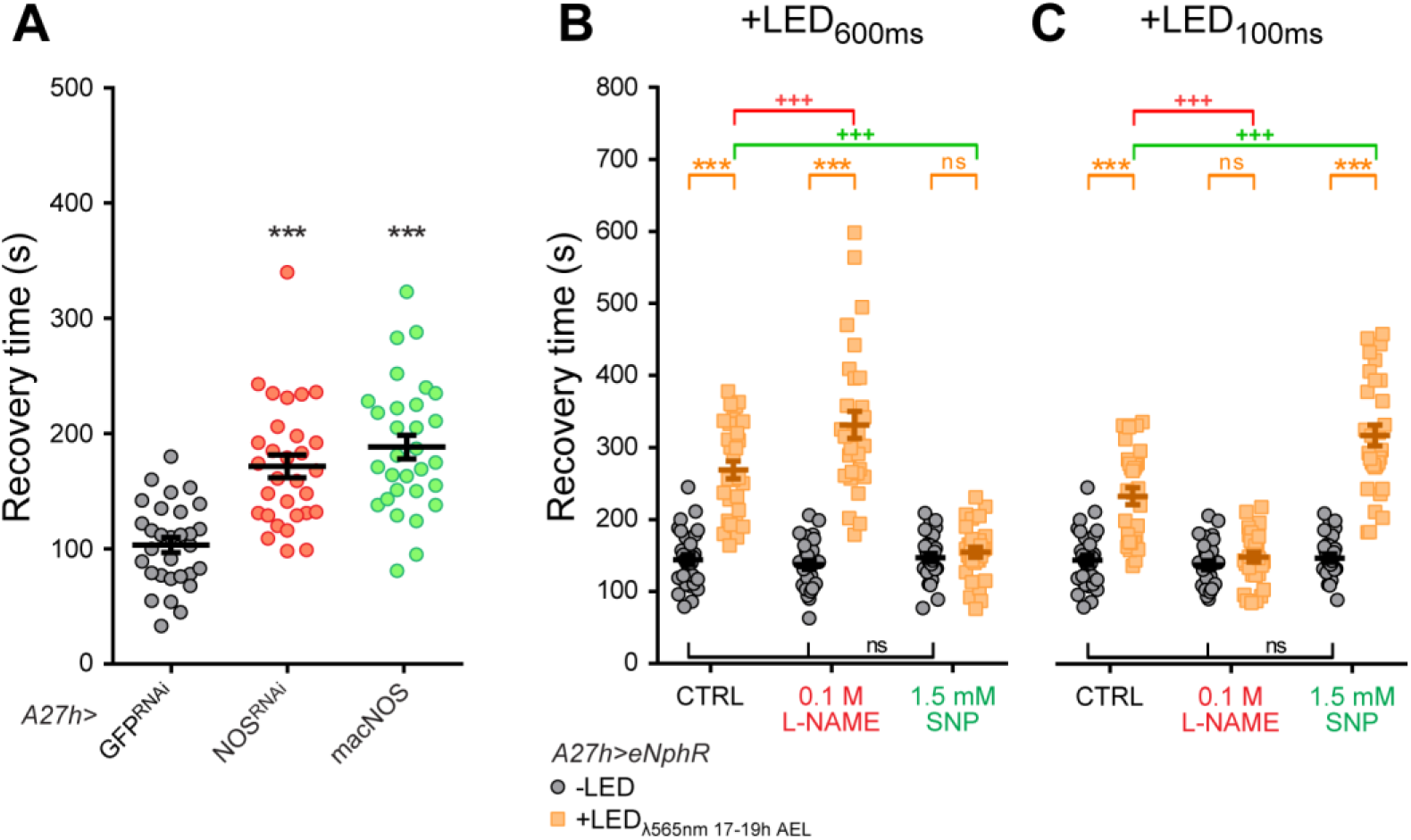
Nitric oxide mediates A27h activity manipulation. (A) Manipulation of NOS in A27h, independently of the direction of change, is sufficient to generate an increase in recovery time to electroshock at L3 (F_(2, 87)_ = 24.74, *p* < 0.0001, One-way ANOVA, followed by Bonferroni’s *post-hoc* test, *n* = 30 in each group, \*\*\**p* < 0.0001). Expression of GFP^RNAi^ was used as a control. (B) Quantitative analysis of recovery time showing the consequence of NO manipulation on A27h inhibition. Embryos, expressing eNpHR in A27h, were exposed to light (λ565 nm, 600 ms – inhibitory stimulation, 17-19h AEL). NOS drugs were administered by feeding gravid females. Pre-exposure to L-NAME significantly potentiated the effect of A27h inhibition, while SNP prevented any increase in recovery time, *n* = 30 in each group. A two-way ANOVA analysis revealed a significant effect of the LED treatment (F_(1, 174)_ = 157.9, *p* < 0.0001), NOS manipulation (F_(2, 174)_ = 32.19, *p* < 0.0001), and interaction (F_(2, 174)_ = 39.65, *p* < 0.0001). (C) Quantitative analysis of recovery time showing the consequence of NO manipulation on A27h excitation. Embryos, expressing eNpHR in A27h, were exposed to light (λ565 nm, 100 ms – excitatory stimulation, 17-19h AEL). The increase in recovery time due to A27h activation was completely prevented by L-NAME or further potentiated by SNP, *n* = 30 in each group. A two-way ANOVA analysis revealed a significant effect of the LED treatment (F_(1, 174)_ = 139.1, *p* < 0.0001), NOS manipulation (F_(2, 174)_ = 45.69, *p* < 0.0001), and interaction (F_(2, 174)_ = 39.13, *p* < 0.0001). (A-C) Data are represented as mean ± SEM. (B, C) \*\*\**p* < 0.001 shows significance to +LED *vs*. -LED within each group; ^+++^*p* < 0.001 shows significance to NOS drugs (+LED groups *vs*. CTRL), Bonferroni’s *post-hoc* test.

Further supporting the involvement of NO in network tuning, embryonic exposure to the NOS-inhibitor, L-NAME (0.1 M) increased the recovery time to electroshock caused by selective inhibition (eNpHR +LED_600ms_) of A27h during the critical period (137 ± 6 *vs*. 331 ± 19 s, -LED *vs*. +LED, respectively, *p* < 0.0001, Fig 5B). Conversely, the presence of the NO-donor, SNP (1.5 mM), blocked the effect of this activity manipulation (147 ± 6 to 154 ± 7 s, -LED *vs*. +LED, respectively, *p* > 0.9). A two-way ANOVA showed that the embryonic manipulation of NO levels significantly affected the optogenetically stimulated (+LED) groups (*p* = 0.0007 and *p* < 0.0001, L-NAME and SNP *vs*. CTRL, respectively), while leaving unstimulated (-LED) animals unchanged, as also shown above. In a complementary set of experiments, inducing hyperactivity in A27h (eNpHR +LED_100ms_) we observed that the same NO-signalling manipulations caused the opposite changes (Fig 5C). Specifically, inhibition of NOS prevented A27h hyperactivity-induced increases in recovery time (0.1 M L-NAME, 137 ± 6 *vs*. 148 ± 7 s, -LED *vs*. +LED, respectively, *p* > 0.9), while exposure to SNP potentiated the increase in recovery time (1.5 mM, 147 ± 6 to 317 ± 15 s, *p* < 0.0001). A two-way ANOVA showed that manipulating NO levels significantly affected the +LED groups only (*p* < 0.0001, both L-NAME and SNP *vs*. CTRL, respectively).

### NO-signalling is sufficient to alter network tuning

Our observation that NO-signalling is necessary for the effect of activity perturbation during the critical period on SRC duration and recovery time in response to electroshock, raises the important question of whether altering NO alone, during the same embryonic period, is sufficient to induce network instability. To this end, we manipulated NO-signalling in all neurons (*elav^C155^*-*Gal4*) in the absence of other activity manipulations; expressing *UAS-NOS^RNAi^* to reduce, or the constitutively active NOS transgene *UAS-macNOS* (22) to increase NO-signalling, respectively. Expression of an RNAi transgene against GFP (*elav^C155^*>*GFP^RNAi^*) was used as control. Both manipulations of NOS were sufficient to significantly increase SRC duration in L3 aCC (NOS^RNAi^: 1.05 ± 0.13 s, *n* = 10, *p* < 0.0001; macNOS: 1.09 ± 0.06 s, *n* = 10, *p* < 0.0001 *vs*. GFP^RNAi^: 0.45 ± 0.02 s, *n* = 13, Fig 6A-B). This mirrors the outcomes achieved by optogenetically increasing or decreasing activity during embryogenesis in otherwise wildtype embryos (5). Genetic manipulation of NO-signalling alone was also sufficient to increase recovery time to electroshock; e.g., pan-neuronal RNAi-mediated knock-down of NOS (*elav^C155^*>*NOS^RNAi^*: 154 ± 8 s, *p* < 0.0001, Fig 6C) or expression of macNOS (*elav^C155^*>*macNOS*: 166 ± 8 s *vs*. CTRL: 101 ± 8 s, *p* < 0.0001).

**Fig 6.**
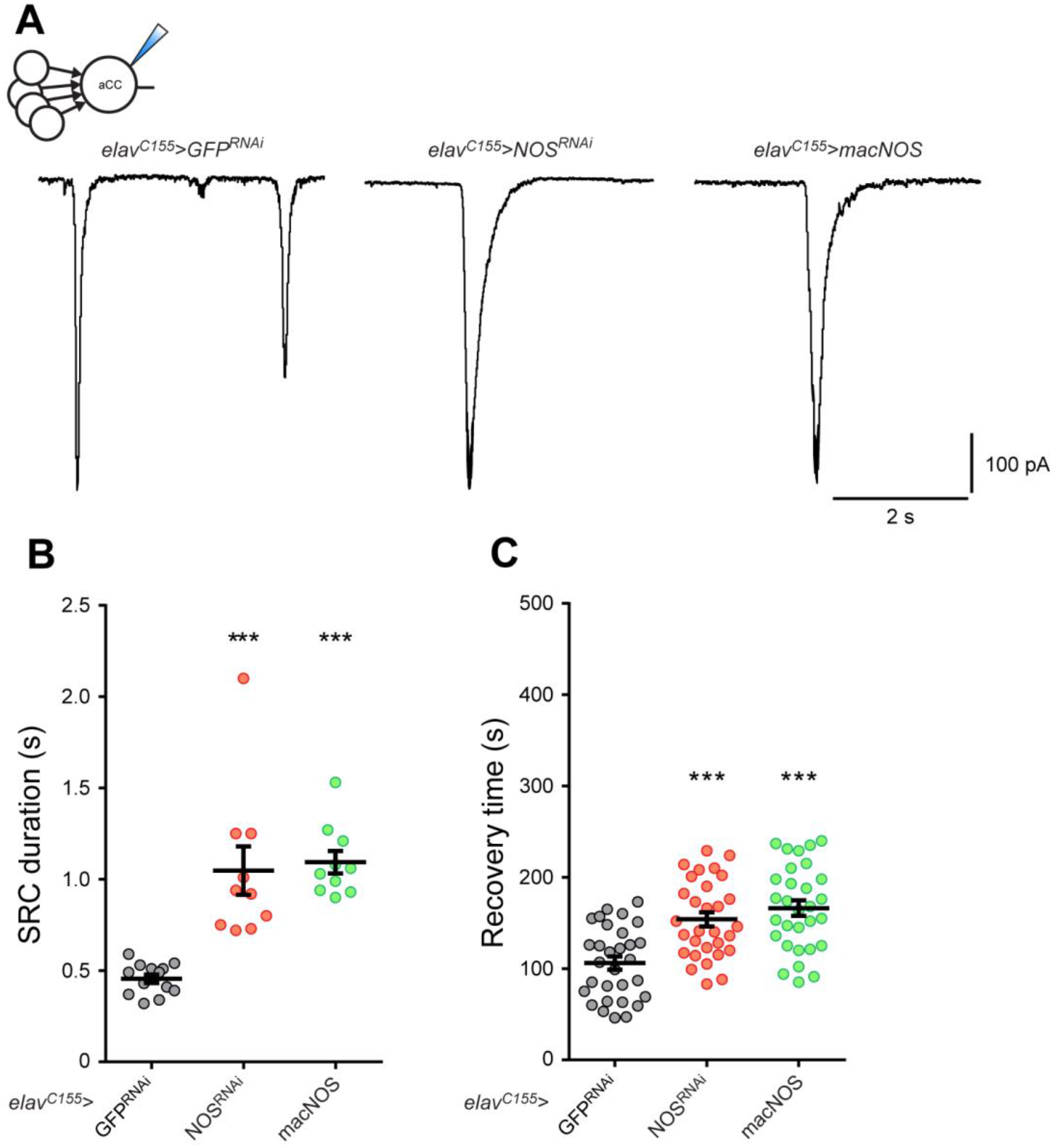
Altered levels of NO during embryogenesis degrade network stability, mimicking optogenetic manipulation. (A) Whole-cell recordings of endogenous SRCs from L3 aCC following pan-neuronal (*elav^C155^*) genetic manipulation of NOS. (B) Pan-neuronal knock down of NOS (NOS^RNAi^, *n* = 10) or macNOS (*n* = 10) overexpression both significantly increased SRC duration. GFP^RNAi^ was used as control (*n* = 13). One-way ANOVA (F_(2, 30)_ = 22.43, *p* < 0.0001) followed by Bonferroni’s *post-hoc* test, \*\*\**p* < 0.0001. (C) Recovery time to electroshock measured from L3 following NOS genetic manipulation. Downregulation of NOS (NOS^RNAi^) and overexpression of macNOS are each sufficient to generate an increase in recovery time. One-way ANOVA (F_(2,87)_ = 16.55, *p* < 0.001) followed by Bonferroni’s *post-hoc* test, *n* = 30 in each group, \*\*\**p* < 0.0001. (B-C) Data are represented as mean ± SEM.

Finally, we observed that exposure of developing embryos to higher concentrations of SNP (5 mM rather than 1.5 mM used previously) is sufficient to induce network instability in L3. Thus, we tested whether activity-manipulation of A27h was sufficient to prevent this. Exposure of gravid females SNP (5 mM) produced L3 with a significantly increased recovery time to electroshock (*A27h*>*eNpHR* + 5 mM SNP -LED: 182 ± 7 s). However, simultaneous inhibition of A27h during the critical period (*A27h*>*eNpHR* + 5 mM SNP +LED_600ms_, 17-19hrs AEL) blocked the effect of SNP (124 ± 5 s, *p* < 0.0001, unpaired *t*-test). This result is consistent with A27h having a significant contribution to network tuning during the critical period.

## Discussion

It is established that activity perturbation during a critical period can induce permanent changes to neural circuit function (3, 4, 23). However, how activity perturbation changes individual synaptic connections across a network and how affected networks respond to such changes is not well understood. In part, this is because our knowledge of critical periods has been largely derived from complex mammalian neural circuits (e.g. vision) that deter single cell resolution. In this respect, it is significant that we have previously identified a critical period during the development of the *Drosophlla* larval locomotor circuitry, during which activity perturbation is sufficient to permanently alter network stability (5). Moreover, we show here that regardless of the nature of the activity manipulation; being inhibitory, excitatory, pharmacological or genetic, the outcome is identical. This predicts that the wiring of this network canalises the impact of activity perturbations, such that they lead to reproducible rather than variable or chaotic outcomes. Moreover, these same outcomes arise regardless of the manipulations having been network-wide or targeted to specific cells. The most parsimonious interpretation for which is that specific components of the motor network (e.g. the pre-motor cholinergic interneuron A27h) have a greater influence on network tuning than others (e.g. the GABAergic A31k interneuron), both when selectively manipulated and also during more global manipulations. Selective A27h activity manipulation is also sufficient to counteract global pharmacological manipulations of NO-signalling which, again, is consistent with a stronger contribution for this neuron to network tuning. The perturbations that we observe are evidently destabilising to mature network function increasing both duration of synaptic currents and severity of induced-seizure activity. Activity disturbance during mammalian critical periods has similarly been linked to detrimental effects: e.g. shift in ocular dominance (e.g. amblyopia), language deficits and autism (4, 24). It has, moreover, been shown that therapeutic intervention to normalise activity during a critical period, in an otherwise activity-disturbed network, can have significant beneficial effects for not only ocular dominance but also epilepsy (4, 5, 25).

Network tuning during a critical period has been studied most extensively in the vertebrate visual system (26–29). However, the underlying mechanisms that have received most attention, for example the role of the homeotic gene OTX2 (30), establishment of perineuronal nets (31) and maturation of GABAergic parvalbumin-positive inhibitory cells (32, 33), are likely not present in *Drosophila*.

Nevertheless, the critical period in the *Drosophila* locomotor circuit coincides with the onset of GABA synthesis (34) and the ingrowth of astrocyte-like glia (35, 36). Whether or not, in *Drosophila*, these events act as instructive cues for the opening and subsequent closure of the critical period remains to be determined. From a circuit perspective in particular, the establishment of an experimental model system such as this, which has an explicit mammalian-like critical period, will greatly facilitate understanding of these enigmatic periods in neural circuit development.

Because neural networks are most likely organised with a hierarchical structure, we reasoned that some constituent neurons may have a greater propensity than others for orchestrating activity-dependent tuning. To validate this hypothesis, we focussed attention on a defined synaptic pairing formed by the excitatory cholinergic premotor interneuron A27h (12) and its postsynaptic motoneuron aCC. Electrophysiology showed that, while pan-neuronal activity manipulations during the critical period lead to overall increased SRCs in aCC motoneurons, a reduction in synaptic strength unexpectedly occurs at the level of this specific pairing. This outcome clearly demonstrates that defined synaptic pairings respond differently to activity perturbation during a critical period. It further demonstrates that neurons that receive multi-neuron synaptic inputs (e.g. aCC) can specifically, and differentially, modify subsets of these inputs. In this example, the A27h premotor IN is differentially affected compared to other excitatory premotor INs to aCC following pan-neuronal activity manipulations during the critical period. Nevertheless, both network-wide (which include A27h) and A27h-specific activity manipulations during the critical period evoke CNS instability. It is significant that the strength of A27h→aCC synaptic transmission is differentially affected by pan-neuronal *vs*. A27h-specific activity manipulation (see Fig 7 for a summary of the effects observed in this study). We cannot, at present, differentiate between whether the changes to network tuning we observe are due to alteration of developmental rules governing the formation of the locomotor circuit or are the consequence of homeostatic changes attempting to maintain stability (37). This discrimination will require more detailed recordings from other neurons of the locomotor circuit, including those that synaptically drive A27h.

**Fig 7.**
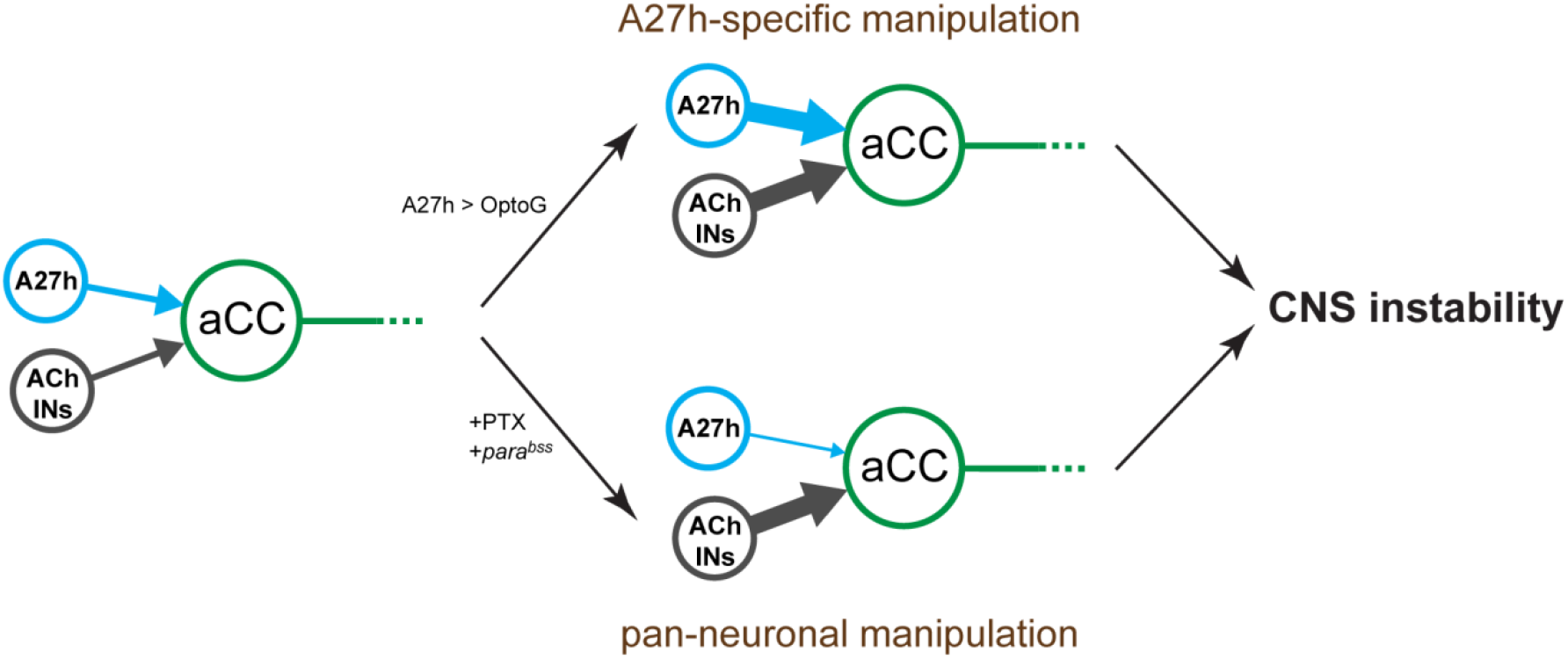
Defined synaptic pairings respond differently to activity perturbation during a critical period. The A27h synaptic drive to aCC is differentially affected by activity perturbation during a critical period. Optogenetic-induced hyperactivity of A27h (specific manipulation) not only potentiates A27h→aCC synapse but also affects other excitatory inputs that aCC receives from the cholinergic premotor INs (ACh INs). Conversely, network-wide perturbation (+PTX or *para^bss^* mutation) reduces A27h→aCC connectivity, though the total aCC inputs are increased. Interestingly, the outcome of both manipulations is CNS instability.

We utilise response to electroshock as a proxy for network stability. A large number of seizure mutations (so-called bang-sensitive mutations) have been identified in *Drosophila* which show seizure-like activity following exposure to a strong stimulus (electroshock or violent shaking) (38).

Similar epilepsy syndromes exist in mammals where exposure to either a loud sound or flashing lights is sufficient to induce a seizure state (39–41). That seizures occur is indicative of underlying networks lacking robustness and which cannot compensate for extremes of activity. At least part of the lack of robustness in *Drosophila* seizure mutants may be due to the increased synchrony seen in activity of motoneurons between adjacent segments (42). Indeed, increased synchrony within neuron subpopulations is a hallmark of mammalian epilepsy (43–45). Thus, whilst we are unable at present to describe the precise effects to global network tuning which results in increased recovery time to electroshock, this measure is an adequate proxy for network robustness which emerges through appropriate network tuning during development.

That single or small subsets of neurons can disproportionately impact the functionality of neural networks, as described in more complex systems, has been described in the mammalian hippocampus and visual cortex (46, 47). In this study, we demonstrate that embryonic manipulation of A27h is sufficient to trigger network instability. Our results also suggest that different interneurons (e.g. A31k *vs*. A27h) may contribute unequally to network tuning, perhaps indicating that their local synaptic connectivity determines how discharges spread from the activated cell to other constituent neurons. Equally, contribution may also be dictated by ability to influence the excitation:inhibition balance of a developing network: in this instance manipulation of excitatory cholinergic neurons (A27h) being more effective than inhibitory GABAergic cells (A31k). Such changes might be expected to differentially perturb the excitation:inhibition balance across a network which may, in turn, contribute to observed postembryonic network stability.

Network tuning requires constituent neurons in any given network to be able to “sense” both their own activity and, importantly, the activity of other neurons within the circuit, regardless of whether they are directly connected. NO, acting as a diffusible signal, is an obvious candidate to fulfil this important signalling role (48–51). Our results support this hypothesis. It is notable that, in this context, we find that either increasing or decreasing NO-signalling generates the same outcome, indicating that change in NO levels, rather than the direction of change, is the key determinant that is crucial for network tuning. The Janus nature of NO has been highlighted in different systems, from cell survival (52), to nociceptive transmission (53) and, particularly, epileptogenesis where its role is highly contradictory with significant evidence supporting both proconvulsive, as well as anticonvulsive activity (54). Thus far, the diversity of the downstream target molecules of NO-signalling, coupled to the lack of homogeneity among these different studies (differing drugs, dose and route of administration applied to diverse seizure models), make it difficult to identify a mechanism. Indeed, it has been reported that NO can differentially modulate excitatory (glutamatergic) and inhibitory (glycinergic) synapses in dorsal horn neurons (53). Our results showing that NO manipulation can produce disparate effects dependent on the endogenous activity level present in the developing circuit may also go some way to explaining these contradictory studies. For example, we show that inhibiting NO-signalling is beneficial to a hyperactive network, but increases network instability (i.e. a proconvulsant action) in a physiologically normal network.

In summary, although a universal phenomenon, how activity tunes a developing network remains poorly understood. We show here that activity perturbation during a defined critical period in the development of the *Drosophila* larval locomotor circuit is sufficient to induce significant and permanent change to network stability. We further show that the effect of activity perturbation is mediated by change to NO-signalling sufficient to induce change to specific synaptic pairings between constituent neurons. The degree of understanding, coupled to an ability to identify and selectively manipulate, the individual neurons that form this circuit offer the prospect of exploiting *Drosophila* to advance understanding of the role of activity in tuning a developing network and, specifically, the role of a critical period in this process.

## Materials and Methods

### *Drosophila* Rearing and Stocks

All *Drosophila melanogaster* strains were grown and maintained on standard corn meal medium at 25°C under a 12:12 h light-dark schedule. Fly stocks include the wildtype strain, Canton S (obtained from Bloomington Drosophila Stock Center, Indiana, USA), and the following previously described transgenic stocks: *UAS-GFP^RNAi^, UAS-NOS^RNAi^, UAS-NOS* and *UAS-macNOS* (a gift of Prof. Oren Schuldiner).

To optogenetically manipulate neurons we used the following lines: channelrhodopsin (55) – *UAS-ChR2_H134R_;cry^03^* (a gift of Dr. Stefan Pulver), halorhodopsin (56) – *UAS-eNpHR::YFP-50C;UAS-eNpHR::YFP-19C, UAS-eNpHR::YFP-34B* (a gift of Prof. Akinao Nose) and *UAS-Chronos* (19).

Expression driver lines include: *elav^C155^-Gal4* – for pan-neuronal expression (57), *elav^C155^-Gal4;cry^03^* – for experiments involving embryonic exposure to blue light, in order to avoid activation of cryptochrome-expressing neurons (58–60). Temporal control of expression was achieved using *Gal80^ts^;OK6-Gal4*, which has been created by crossing *tub*>*Gal80^ts^* and *OK6-Gal4* – for expression in motoneurons (61), (both obtained from Bloomington Drosophila Stock Center). *A27h-Gal4* (*R36G02-Gal4*) – for selective expression in the premotor interneuron A27h (12). This line was also crossed with the mutant *bang senseless^1^* to drive transgenes in a bang sensitive background (*para^bss^;; A27h-Gal4*). Finally, the *A31k split-Gal4* line (*20A03-AD;87H09-DBD*, a gift of Prof. Chris Doe) – for selective expression in the GABAergic interneuron A31k (11).

A27h and A31K driver lines were tested during embryogenesis for the expression of a fluorescent UAS-reporter, *UAS-CD4::tdTomato* (62). Confocal acquisitions confirmed reliable *CD4::tdTomato* expression during the critical period (17-19 h AEL).

### Optogenetic manipulation of neuronal activity

Mated adult females were allowed to lay eggs on grape agar (Dutscher, Essex, UK) plates at 25°C supplemented with a small amount of live yeast paste. To ensure that embryos received enough retinal, adults were fed with 4 mM *all-trans*-retinal (Sigma-Aldrich, Poole, UK) dissolved in yeast paste twice a day for three days prior to collection. Embryos were collected within a 4 h time range (time 0±2 h after egg laying) and then transferred to a fresh grape agar plate. Plates were placed in a humidified atmosphere inside a 25°C incubator and exposed to collimated light from an overhead LED, positioned 17 cm from the embryos. LEDs had peak emission at λ470 nm (bandwidth 25 nm, irradiance 466 ± 14 nW·cm^-2^; OptoLED, Cairn Instruments, Kent, UK) or λ565 nm (bandwidth 80 nm, 250 ± 10 μW·cm^-2^; M565L2, Thorlabs, Newton, NJ). Embryonic exposure to λ470 nm, but not λ565 nm is sufficient to induce a seizure-phenotype at L3 (5) through the activation of cryptochrome-expressing neurons (58–60). Therefore, experiments involving blue light required a cry-null (*cry^03^*) background, to avoid unspecific effects. Alternatively, we used either eNphR or Chronos, a green-shifted variant of ChR, which allowed us to activate specific subpopulations of neurons during embryogenesis without simultaneous activation of the endogenous blue-light sensitive cry neurons.

Light was pulsed at 1 Hz using a Grass S48 stimulator (Grass instruments, Quincy, MA, USA). ChR and Chronos were activated with short duration pulses (100 ms). In neurons overexpressing eNpHR, the duration of light pulses varied from 100 ms to 600 ms in order to induce rebound firing activity (hyperactivity) or preventing spike firing (inhibition), as previously described (5).

Embryos were optically treated for a pre-determined time period during embryogenesis (between 17-19^th^ ± 2 h AEL), which corresponds to the critical period (5). After manipulation, embryos were transferred into food bottles and maintained at 25°C in complete darkness until ~4 days later when wall-climbing L3 were collected and then electrophysiologically or behaviourally tested.

### Drug Treatments

Embryonic exposure to Picrotoxin (PTX, Sigma-Aldrich) was achieved by feeding gravid females to live yeast paste supplemented with 0.25 mg/ml PTX for three days, prior to embryo collection. Embryos were collected as previously described and transferred to nondrug-containing vials (5).

Chemical manipulation of the NO pathway was performed using the same procedure. Feeding gravid females low doses of L-NAME (0.1 M, Sigma-Aldrich), or SNP (1.5 mM, Sigma-Aldrich), was sufficient to affect the outcome of the optogenetic stimulation (increased recovery time and prolonged SRC duration) whilst leaving the control group (-LED) unaltered. Conversely, higher doses (0.5 M L-NAME or 5 mM SNP), although more effective, were sufficient to alone mimic the effect of optical stimulation (ChR -LED or in a WT strain, Canton S) thus affecting control levels. We obtained identical results by testing other inhibitors/activators targeting the NO pathway. Hence, for all the experiments shown in this paper, we first pre-determined an optimal concentration for each compound by testing a wide range of doses (data not shown).

### Quantitative RT-PCR

Total RNA was extracted from 10 L3 CNSs using the RNeasy^®^ mini kit (#74104 QIAGEN, West Sussex, UK). cDNA synthesis was carried out in 20 μl total volume using the RevertAid H Minus First Strand cDNA Synthesis Kit (#K1632, Thermo Fisher Scientific, Loughborough, UK). Oligo(dT) (0.5 μg) and random hexamers (0.2 μg) were mixed with RNA and made up to 12 μl with RNase-free water. The mix was incubated at 65°C for 5 min to denature RNA followed by incubation on ice for 1 min. To this was added 4 μl of reaction buffer (in mM: 250 Tris-HCl, 250 KCl, 20 MgCl_2_, 50 DTT), 2 μl of 10 mM dNTPs, 1 μl of RNase inhibitor (Ribolock R1) and 1 μl of RevertAid H Minus reverse transcriptase. The reaction was incubated at 25°C for 10 min, 42°C for 60 min followed by 70°C for 10 min.

Quantitative PCR was performed using SYBR Green I real-time PCR method (LightCycler^®^ 480 SYBR^®^ Green I Master, Roche, West Sussex, UK). The Ct values, as defined by the default setting, were measured using a LightCycler^®^ 480 II real-time PCR (Roche) using a thermal profile of 10 min at 95°C followed by 45 cycles of 10 s at 95°C, 10 s at 60°C and 10 s at 72°C. Single-product amplification was confirmed by post-reaction dissociation analysis. PCR primers against the *dNOS* mRNA sequence were designed with the aid of LightCycler^®^ Probe Design Software 2.0 (v1.0) (Roche). NOS Primer sequences (5’ to 3’) are: forward GCAGGAATTCGATTCGTTGT and reverse GATGCCAAAGATGTCCTCGT. For each biological sample, three technical replicates were tested for the gene of interest as well as for the reference gene (actin). Relative gene expression was calculated as the 2^−ΔCt^, where ΔCt was determined by subtracting the average actin Ct value from that for each gene.

The efficacy of the *UAS-NOS^RNAi^* and *UAS-dNOS* transgenes, pan-neuronally co-overexpressed with channelrhodopsin (*elav^C155^*>*ChR;NOS^RNAi^* and *elav^C155^*>*ChR;dNOS*, respectively), was tested by quantitative real-time PCR. Both manipulations showed a significant changes in *dNOS* mRNA: NOS^RNAi^ = −1.11 ± 0.25 fold change, *p* = 0.0154; NOS = +5.61 ± 1.48, *p* = 0.0039; one-way ANOVA (F_(2,15)_ = 16.73, *p* < 0.0002) followed by Bonferroni’s *post-hoc* test, *n* = 6 in each group.

### Electrophysiology

Electrophysiological recordings were performed in L3 as previously described (6, 18). Whole cell voltage- and current-clamp recordings were achieved using thick-walled borosilicate glass electrodes (GC100F-10, Harvard Apparatus, Edenbridge, UK) fire polished to resistances of 10-15 MΩ. Recordings were made using a Multiclamp 700B amplifier controlled by pCLAMP (version 10.4) via a Digidata 1440A analog-to-digital converter (Molecular Devices, Sunnyvale, CA). Traces were sampled at 20 kHz and filtered online at 10 kHz. External saline composition was as follows: 135 mM NaCl, 5 mM KCl, 4 mM MgCl_2_·6H_2_O, 2 mM CaCl_2_·2H_2_O, 5 mM TES and 36 mM sucrose, pH 7.15. Internal patch solution was as follows: 140 mM K^+^-D-gluconate, 2 mM MgCl_2_·6H_2_O, 2 mM EGTA, 5 mM KCl, and 20 mM HEPES, pH 7.4. KCl, CaCl_2_, MgCl_2_ and sucrose were purchased from Fisher Scientific (Loughborough, UK); all remaining chemicals were obtained from Sigma-Aldrich (Poole, UK).

A27h→aCC synaptic drive was measured by stimulating A27h interneurons expressing ChR and recording the postsynaptic response from aCC motoneurons, in whole-cell configuration, at the same time. ChR was activated with a λ470 nm LED (OptoLED, Cairn Instruments, Kent, UK) connected to an Olympus BX51WI microscope. During recording, light was pulsed onto the sample for 1 s triggered by TTL signals from pClamp (Molecular Devices) to the LED controller. The same stimulation protocol was applied five times to each neuron and the recordings averaged. Current amplitudes were measured as the maximal current evoked and normalised for cell capacitance.

Spontaneous rhythmic currents (SRCs) were recorded from L3 aCC motoneurons for 3 minutes. Traces were sampled at 20 kHz and filtered at 0.2 kHz low pass. Cells with input resistance <0.5 GΩ were not considered for analysis. Synaptic current parameters were examined for each recorded cell using Clampfit (version 10.4). To measure the amplitude of SRCs, the change from baseline to peak current amplitude was determined (63). Currents shown were normalized for cell capacitance (determined by integrating the area under the capacity transient resulting from a step protocol from −60 to −90 mV). The duration of each synaptic event was defined as the time from current initiation until the return to baseline, as depicted in Fig 1B.

Loose patch recordings were performed on L3 aCC motoneurons. Thin-wall borosilicate glass capillaries (GC100TF-10, Harvard Apparatus) were used to pull recording electrodes (unpolished) with resistances between 1.5–2.5 MΩ. Data were acquired with a sampling rate of 20 kHz, filtered with a low-pass filter of 0.2 kHz and analysed in Clampfit 10.4 (Molecular Devices). Excitation of A27h expressing Chronos was achieved using an OptoLED system (Cairn Research) with a 565 nm LED. For each cell, the contribution of A27h to aCC spiking activity was quantified by delivering light pulses of 1 second at 0.2 Hz and averaging the number of action potentials fired in the first 10 pulses. Bouts overlapping with endogenous activity were rejected.

### Electroshock assay

Electroshock assay was performed as previously described (18). Briefly, wall-climbing L3 were transferred to a plastic dish after washing to remove food residue and gently dried using paper tissue. Once normal crawling behaviour resumed, a conductive probe, composed of two tungsten wires (0.1 mm diameter, ~1-2 mm apart) was positioned over the approximate position of the CNS, on the anterior-dorsal cuticle of the animal. A 2.3 V DC pulse for 2 s, created by a constant voltage generator (DS2A-mkII, Digitimer Ltd., Welwyn Garden City, Hertfordshire, UK), was applied. In response to the stimulus, we observed a transitory paralysis in which larvae were tonically contracted and, occasionally, exhibited spasms. The time to resumption of normal crawling behaviour was measured as recovery time. Normal crawling was defined as a whole body peristaltic wave resulting in forward movement.

### Quantification and Statistical Analysis

Data was acquired and imported into Microsoft Excel (Microsoft Corp., Redmond, WA). All statistical tests were performed in GraphPad Prism (version 7). All data are presented as the mean ± SEM. Details including the exact value of *n* for each sample and *P* values are provided in the Results section or in each respective figure legend.

Statistical analyses were conducted using either unpaired *t*-test (Fig 1D, 2C-D), one-way (Fig 2E-G, 3B, 3D, 3F 4D-F, 5A-B, 6B-C, and Fig S1), or two-way (Fig 4B-C, 5B-C) ANOVA as indicated in the respective figure legends. For one-way or two-way ANOVAs, *post-hoc* Bonferroni’s multiple comparisons tests were conducted. Significance was shown as * = *p* < 0.05, ** = *p* < 0.01, *** = *p* < 0.001, and not significant values were not noted or shown as ns. Figures were assembled with Adobe Illustrator CS3 (Adobe, San Jose, CA, USA).

## Acknowledgements

The authors would like to thank Oren Schuldiner for *UAS-macNOS* flies, Stefan Pulver for *UAS-ChR2_H134R_* flies, Akinao Nose for *UAS-eNpHR*, and Chris Doe for the A31k split-Gal4 line. This work was supported by funding from the Biotechnology and Biological Sciences Research Council to R.A.B. (BB/N/014561/1) and to M.L. (BB/R016666/1). Work on this project benefited from the Manchester Fly Facility, established through funds from the University and the Wellcome Trust (087742/Z/08/Z).

## Author Contributions

C.N.G.G. and R.A.B. designed experiments. C.N.G.G. and Y.N.F. performed experiments and analysed data. M.L. verified correct expression of driver lines. C.N.G.G., M.L. and R.A.B. wrote the paper.

---

**S1 Fig.**
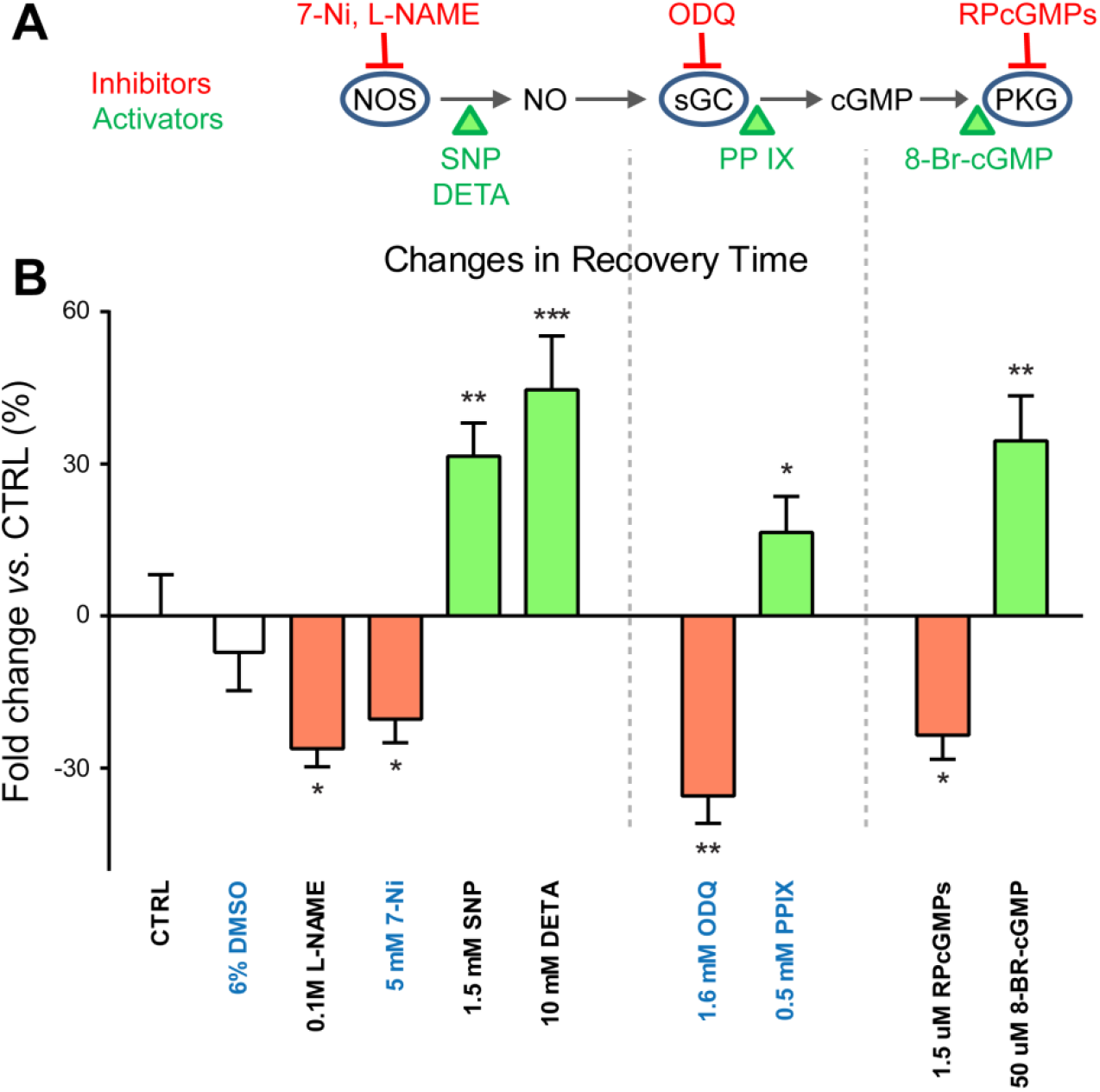
The canonical nitric oxide signalling pathway mediates activity perturbation during the critical period. (A) Schematic representation of the NO-signalling pathway. NOS: Nitric oxide synthase, sGC: soluble guanylyl cyclase, cGMP: cyclic guanosine monophosphate and PKG: Protein Kinase G. Inhibitors and activators were indicated in red and green, respectively. (B) Chemical manipulation of the NO pathway affects the ChR-induced increase in recovery time to electroshock (*elav^C155^*>*ChR*, LED_100ms_, 17-19h AEL). Values are expressed as fold change (+LED/-LED) and compared to the control (set to zero). For each compound, drug concentration was first optimised to ensure no affect was observed in the -LED control group (One-way ANOVA F_(9, 290)_ = 0.8452, *p* = 0.4987). Drugs labelled in blue were dissolved in 6% DMSO (which had no effect: −6.9 ± 7.9%). All inhibitors significantly reduced the ChR-increase in recovery time (0.1 M L-NAME: −26.2 ± 3.6%, *p* = 0.016; 5 mM 7-Ni: −20.4 ± 4.6%, *p* = 0.041; 1.6 mM ODQ: −36.53 ± 5.4%, *p* = 0.005 and 1.5 μM RPcGMPs: −23.52 ± 4.7%, *p* = 0.018), while activators potentiated the effect of ChR activation (1.5 mM SNP: +31.5 ± 6.6%, *p* = 0.0048; 10 mM DETA: +44.6 ± 10.6%, *p* = 0.0002; 0.5 mM PPIX: +16.5 ± 7.1%, *p* = 0.0341 and 50 μM 8-BR-cGMP: +34.5 ± 8.9%, *p* = 0.0040). One-way ANOVA (F_(9, 290)_ = 16.86, *p* < 0.001) followed by Bonferroni’s *post-hoc* test, *n* = 30 in each group. Data are represented as mean ± SEM.

